# Biosynthesis of Circular RNA ciRS-7/CDR1as Is Mediated by Mammalian-Wide Interspersed Repeats (MIRs)

**DOI:** 10.1101/411231

**Authors:** Rei Yoshimoto, Karim Rahimi, Thomas Hansen, Jørgen Kjems, Akila Mayeda

**Affiliations:** Division of Gene Expression Mechanism, Institute for Comprehensive Medical Science, Fujita Health University, Toyoake, Aichi, 470-1192, Japan; Department of Molecular Biology and Genetics, Aarhus University, C.F. Møllers Allé 3, 8000 Aarhus C, Denmark; Interdisciplinary Nanoscience Center, Aarhus University, Gustav Wieds Vej 14, 8000 Aarhus C, Denmark

## Abstract

Circular RNAs (circRNAs) are stable noncoding RNAs with a closed circular structure. One of the first and best studied circRNAs is ciRS-7 (CDR1as) that acts as a regulator of the microRNA miR-7, however, the biosynthesis pathway has remained an enigma. Here we delineate the biosynthesis pathway of ciRS-7. The back-splicing events that form circRNAs are often facilitated by flanking inverted repeats of the primate-specific *Alu* elements. ciRS-7 gene lacks these elements but, instead, we identified a set of flanking inverted elements belonging to the mammalian-wide interspersed repeat (MIR) family. Splicing reporter assays in HEK293 cells demonstrated that these inverted MIRs are required to generate ciRS-7 through a back-splicing and CRISPR/Cas9-mediated deletions confirmed the requirement of the endogenous MIR elements in SH-SY5Y cells. Using bioinformatics searches, we identified several other MIR-dependent circRNAs that we confirmed experimentally. We propose that MIR-mediated RNA circularization constitutes a new widespread biosynthesis principle for mammalian circRNAs.

## INTRODUCTION

Endogenous circular RNAs (circRNAs) were first observed as scrambled exon transcripts (Nigro et al., 1991) and these transcripts were found to have circular structures with covalently closed ends (Capel et al., 1993; Cocquerelle et al., 1993). For two decades, however, they were disregarded as rare oddities, or regarded as poorly expressed mis-spliced products.

High-throughput sequencing of full transcriptomes (RNA-Seq) has identified thousands of circRNAs in eukaryotes and these are now considered common byproducts of many protein-coding genes (Salzman et al., 2012; Jeck et al., 2013; Memczak et al., 2013). It has been reported that circRNA generation *via* back-splicing and authentic mRNA production *via* conventional splicing from the same pre-mRNA are mutually exclusive events, suggesting that circRNA formation potentially regulates the expression of the host gene (Ashwal-Fluss et al., 2014; reviewed in Wilusz, 2018).

The functions of the vast majority of circRNAs remain unknown, however, some of these circRNAs have important biological functions. For instance, certain circRNAs control the stability and activity of micro RNAs (miRNAs), regulate transcription or alternative splicing, affect translation of host genes, or they can even be translated and produce proteins themselves (reviewed in Wilusz, 2018). ciRS-7 (also known as CDR1as) was one of the first functionally annotated circular RNAs. It is conserved among mammals and is mainly expressed in the brain. ciRS-7 has many binding sites for a particular miRNA, miR-7, and a single binding site for miR-671 that triggers Argonaute2-catalized slicing of ciRS-7 (Hansen et al., 2011). *In cellulo* experiments suggested that ciRS-7 may function as a sponge, or decoy, that reduces the available free miR-7 and thus prevents repression of miR-7 targeted mRNAs (Hansen et al., 2013; Memczak et al., 2013). Knockout mice lacking the ciRS-7 genomic locus down-regulated miR-7 in the brain, suggesting that ciRS-7 has a role in stabilizing or transporting miR-7 (Piwecka et al., 2017). As a result, the ciRS-7 knockout mice had impaired sensorimotor gating, which is associated with neuropsychiatric disorders.

Recent gene editing experiments revealed a comprehensive regulatory network in mouse brain between a long non-coding RNA (lncRNA), circRNA, and two miRNAs (Kleaveland et al., 2018). This network involves the Cyrano lncRNA that promotes the degradation of miR-7, which in turn enhances the miR-671-directed degradation of ciRS-7, meaning that Cyrano causes an accumulation of ciRS-7. ciRS-7 is apparently a key component in a gene-regulatory network in the brain but understanding the biosynthesis and transport of this particular circRNA remains an important challenge.

It is particularly important to understand the biosynthesis of endogenous ciRS-7. In an artificial ciRS-7 expression vector, the insertion of 800 nucleotides (nt) of perfectly complementary sequences into an intron is technically sufficient to circularize ciRS-7 (Hansen et al., 2013). Evidently there is no such extensive stretches of complementarity nearby the endogenous ciRS-7 exon. In human, the pairing of inverted repeats of *Alu*, a primate-specific repetitive element, has been claimed to promote direct back-splicing in a subgroup of circRNAs (Jeck et al., 2013; Liang and Wilusz, 2014; Zhang et al., 2014; Venø et al., 2015; Zheng et al., 2016). Here we showed that highly conserved mammalian-wide interspersed repeat (MIR) sequences, but not *Alu* sequences, in the flanking introns of the ciRS-7 are required for back-splicing to generate the circular RNA structure. Bioinformatics analyses followed by reporter assays furthermore identified additional distinct circRNAs generated by inverted MIR sequences. Here we demonstrate that a subset of mammalian circRNAs are generated by mammalian-wide MIR-mediated back-splicing.

## RESULTS

### Experimental Evidence Supports ciRS-7 Generation through Back-Splicing

Two major circRNA biosynthesis pathways have been proposed (Figure S1A; reviewed in Jeck and Sharpless, 2014; Wilusz, 2018). Both pathways involve a splicing reaction between downstream 5’ and upstream 3’ splice sites of the circularized exon(s), however, this splicing event occurs either directly on the loop-structure formed *via* the flanking intronic complementary sequences in a ‘Back-splicing pathway’, or on the lariat-structure generated by exon-skipping splicing in an ‘Intra-lariat splicing pathway’.

To determine the pathway used to generate ciRS-7, we first examined the ciRS-7 precursor transcript (~80 kb) that includes six exons and five introns (Figure S2). The last exon contains the entire ciRS-7 sequence and we identified new flanking alternative 5’ and 3’ splice sites (Figure S2). Our splice site targeting experiments with antisense oligoribonucleotides (ASOs) suggested that the ‘Back-splicing pathway’, but not ‘Intra-lariat pathway’, produces circular ciRS-7 (Figure S1B, C, D). This finding was further validated by a genomic deletion of these relevant splice sites by CRISPR/Cas9-mediated technology (Figures S4).

### The ciRS-7 Locus Is Flanked by Inverted MIR Sequences

Since our results suggested that the ‘Back-splicing pathway’ but not the ‘Intra-lariat splicing pathway’, is responsible for generating ciRS-7, we first searched for the inverted *Alu* elements thought to be required for the ‘Back splicing pathway’. Using the UCSC RepeatMasker (in human hg19 coordinates) we identified two human *Alu* elements upstream of ciRS-7 within exon 5 and exon 6 (Figure S3), but found no downstream *Alu* element that could generate the required base-pairing around the ciRS-7 exon. Therefore, flanking inverted *Alu* elements are unlikely to account for the generation of ciRS-7.

However, we did find two relatively conserved regions upstream and downstream of the ciRS-7 exon in an inverted orientation that are conserved both in human and mouse (Figure 1A). These regions overlapped with mammalian-wide interspersed repeats (MIRs), an ancient short interspersed nuclear elements (SINE) family conserved among mammals and marsupials (Jurka et al., 1995; Krull et al., 2007). These findings suggest that ciRS-7 could be generated *via* a back-splicing pathway facilitated by inverted MIRs.

**Figure 1.**
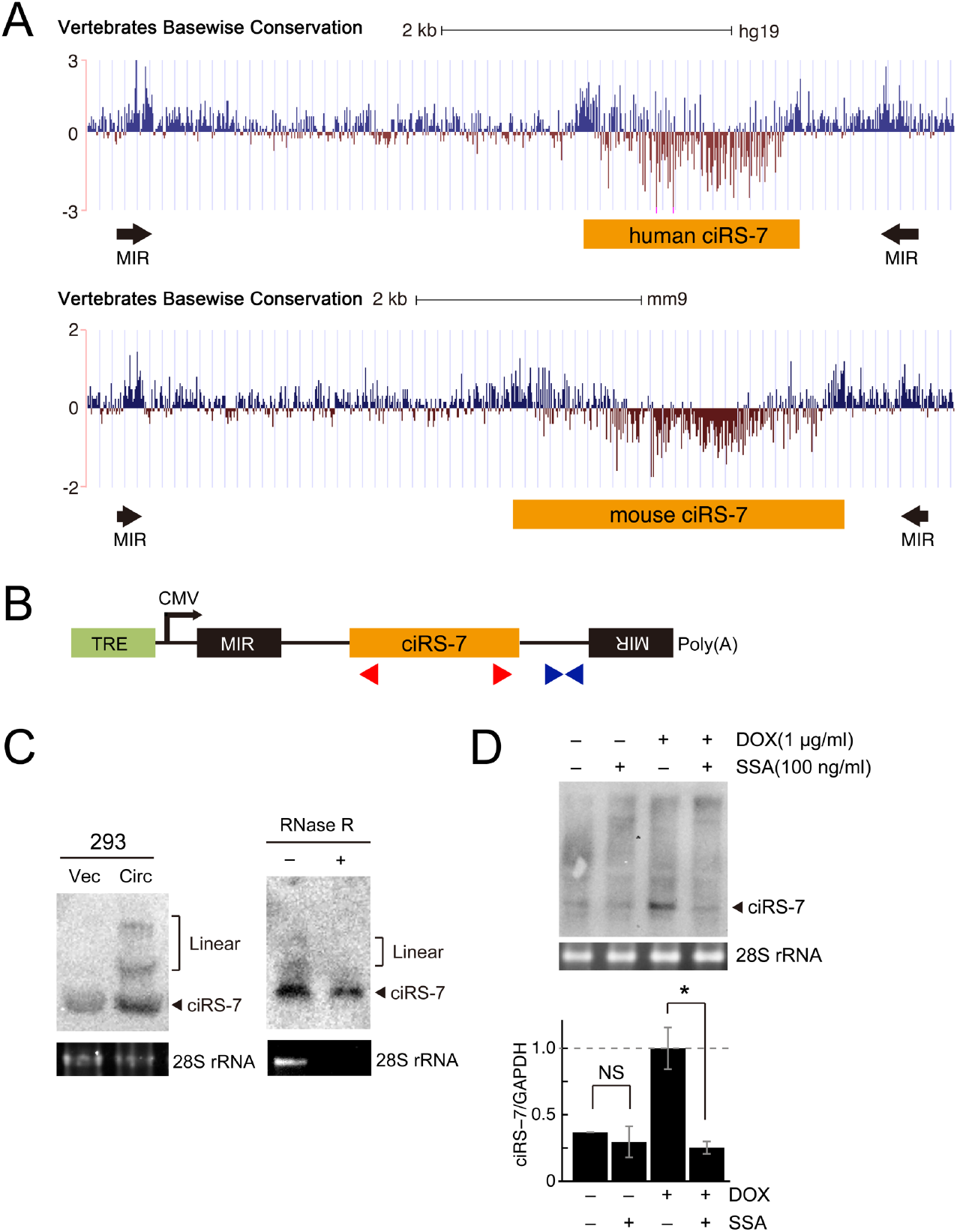
The Production of ciRS-7 Depends on the Flanking Inverted Sequences of MIRs. **(A)** The ciRS-7 locus (human/mouse ciRS-7) and the identified conserved inverted MIR elements (MIRs) in the genome browser track. The vertebrate basewise conservations of human (upper) and mouse (lower) genomic sequences around the ciRS-7 precursor were displayed in the UCSC Genome Browser. **(B)** Schematic structure of ciRS-7-expressing reporter plasmids. Transcription is driven by a CMV promoter and an arrow denotes the initiation site. The tetracycline response element (TRE), inverted MIR elements, ciRS-7 coding exon (ciRS-7) are indicated. Red and blue triangles indicate the positions of PCR primers to detect ciRS-7 and its precursor, respectively. **(C)** The generation of ciRS-7 from reporter in stably transduced HEK293 cells. The cells were treated with DOX and extracted total RNA was analyzed by Northern blotting (left panel). 28S rRNA was visualized by ethidium bromide staining as a loading control. To validate the circular structure of generated ciRS-7, total RNA was also treated with RNase R prior to the Northern blotting analysis (right panel). **(D)** The effect of splicing inhibitor, SSA, on the production of ciRS-7. The same stably transduced cells described in panel C were treated with DOX (+ lanes) followed by SSA addition (+ lanes). After 12 h culture, extracted total RNA was analyzed by Northern blotting to detect ciRS-7 (upper panel). Ethidium bromide-stained 28S rRNA was shown as a loading control. The total RNA was also analyzed by RT–PCR and the graph shows quantification of the ciRS-7 generation (lower panel). The ciRS-7 expression levels were normalized to the control expression level of GAPDH and plotted as ratios to the value of DOX induced cells. Means ± SD are given for two independent experiments (*P < 0.05, NS=not significant).

### Inverted MIRs Promote ciRS-7 Biosynthesis

To test this hypothesis, we generated a stable HEK293 cell line that expresses a doxycycline (DOX)-inducible 5-kb transcript containing the ciRS-7 exon flanked by upstream and downstream inverted MIRs (Figure 1B).

The ciRS-7 circular product and its precursor RNA were both detected by Northern blot 24 h after induction with DOX (Figure 1C, left panel). We verified that the final ciRS-7 product was indeed circular by pre-treatment with RNase R (Figure 1C, right panel). These results demonstrate that our mini-gene, covering the ciRS-7 exon and its inverted MIRs, recapitulates the endogenous generation of circular ciRS-7. To test if ciRS-7 generation from the mini-gene system is splicing-dependent, we used the general pre-mRNA splicing inhibitor Spliceostatin A (SSA) (Kaida et al., 2007; Yoshimoto et al., 2017). SSA clearly inhibited DOX-induced ectopic ciRS-7 production, suggesting that ciRS-7 is indeed generated by splicing (+DOX lanes, Figure 1D).

Using this stable cell line system, we next made a series of MIR-deleted mini-genes to analyze the role of MIRs in human ciRS-7 biosynthesis (Figure 2A). The wild-type mini-gene efficiently generated ciRS-7 (Figure 2B; WT), whereas the deletion of either the upstream or downstream MIR element seriously impaired ciRS-7 production (Figure 2B; Δ5’MIR, Δ3’MIR, Δ5’3’MIR), demonstrating that ciRS-7 generation depends on both inverted MIRs.

**Figure 2.**
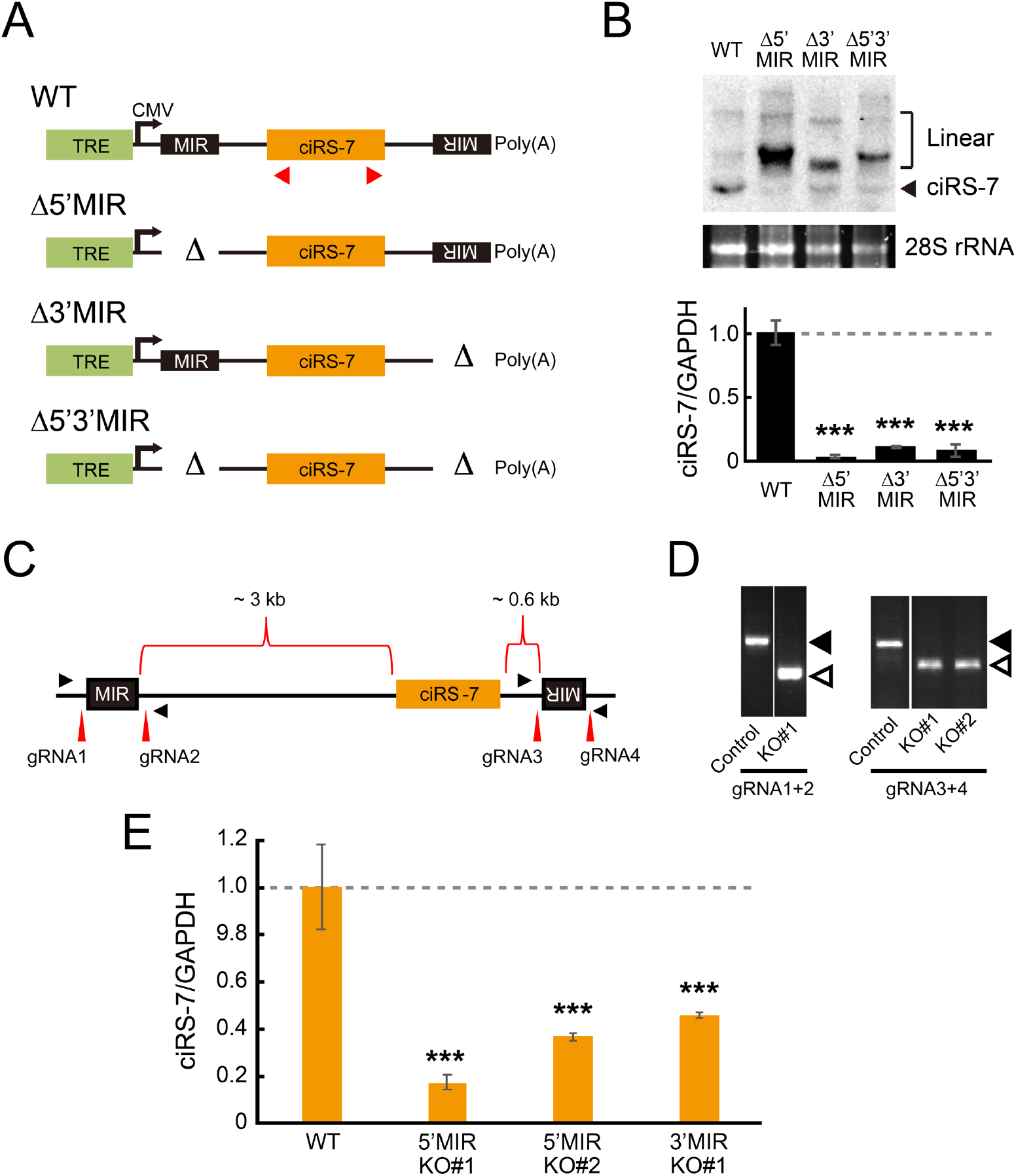
Deletion of Either Upstream or Downstream MIRs Aborts the Production of ciRS-7. **(A)** Schematic structure of ciRS-7-expressing reporters with or without MIR sequence (see Figure 2B for the abbreviations). Red triangles indicate the positions of PCR primers to detect ciRS-7 in panel B. **(B)** Generation of ciRS-7 from reporters, with or without MIR sequences, in stably transduced HEK293 cells. Total RNA extracted from DOX-treated cells was analyzed by Northern blotting (upper panel). 28S rRNA was detected by ethidium bromide staining as a control. Total RNA was also analyzed by RT–PCR and the graph shows quantification of the ciRS-7 generation (lower panel). The ciRS-7 expression levels were normalized to the control expression level of GAPDH and plotted as ratios to the value of control wild-type (WT) plasmid-expressing cells. Means ± SD are given for three independent experiments (***P < 0.001). **(C)** Schematic genomic structure of ciRS-7 locus with the flanking inverted MIR sequences. The positions of guide RNAs (gRNA1–gRNA4) to delete each MIR element are indicated with red vertical arrowheads. PCR primers for detecting deleted sites are indicated with filled triangles. **(D)** The genomic deletions of the flanking MIR elements in SH-SY5Y cell clones were verified by genomic PCR. The indicated two pairs of gRNAs were used to delete the 5’ and 3’ MIR elements. PCR primers indicated in panel A were used for detecting the MIR deletion. Open and filled triangles point to the deleted (KO#1, KO#2) and non-deleted (Control) alleles, respectively. **(E)** The effect of the homozygous MIR deletions (5’MIR KO and 3’MIR KO) on ciRS-7 production was quantified by RT–qPCR. The ciRS-7 expression levels were normalized to the control expression level of GAPDH in the same clones (ciRS-7/GAPDH). Values are relative to the value of control wild-type clone (WT). Means ± standard deviation (SD) are given for three independent experiments (***P < 0.001).

Finally, we used CRISPR/Cas9-mediated editing to delete the endogenous MIR sequences in the neuronal SH-SY5Y cell line. Two guide RNAs (gRNAs) sets were designed to target the boundaries of the 5’ and 3’-MIR sequences (Figure 2C) and co-transfected with the Cas9 expression vector to delete the MIRs. The efficiencies of targeted genomic deletion were determined by PCR analyses using primers flanking the targeted regions and the PCR-amplified fragments were resolved by gel electrophoresis (Figure 2D). We observed that circularization was markedly inhibited when either the upstream or downstream MIR sequence was deleted (Figure 2E). Together, we conclude that the ciRS-7 is generated *via* the back-splicing pathway promoted mainly by its flanking inverted MIR sequences.

### Other MIR-Dependent and MIR-Independent circRNAs Were Identified

Since MIRs are ancient SINEs and ubiquitous in mammalian genomes (Jurka et al., 1995), we speculated that the utilization of MIRs in promoting back-splicing to generate ‘MIR-dependent’ circRNAs could be a globally conserved strategy rather than a gene-specific event.

Upon bioinformatics analysis, we indeed found that MIRs neighboring ciRS-7 are conserved among Primates, Euarchontoglires, Laurasiatheria, Afrotheria, and Armadillo. Here we show the maps of ciRS-7 with the MIRs of human and mouse (Figure 1A). We went on to search for other potential MIR-dependent circRNAs. Using the RepeatMasker track in human hg19 coordinates, we found a total of 595,094 MIRs. Comparing with the mouse mm9 coordinates, 104,074 are conserved between human and mouse. Previously, 7,771 circRNAs were identified from RNA-Seq data of foreskin cells (Jeck et al., 2013). Using this data set, we identified 170 circRNA candidates that contain inverted intronic MIRs within 1,000 nt upstream and downstream of the circularized exons.

From these 171 circRNA candidates, we first eliminated circRNAs with very large precursors (>5 kb) since these are hard to validate using mini-gene reporters. Among the circRNAs showing ubiquitous expression, we arbitrarily chose five circRNAs as representatives; circCDK8, circSPNS1, circTMEM109, circZYMND8, and circSRGAP2C (Figure S5 for the maps with identified MIRs). To test whether biosynthesis of these circRNAs is MIR-dependent or not, we made a series of MIR-deleted mini-genes as we had done for ciRS-7 (Figure 3A). Plasmids expressing these mini-genes were transfected into mouse N2A cells, in which ectopically expressed human circRNAs could be discriminated from the endogenously expressed mouse circRNAs. All five circRNAs were successfully detected from the wild-type mini-genes and their circular structure was verified by RNase R digestion (Figure 3B). Deletion of either upstream or downstream MIRs prevented the production of circCDK8 and circSPNS1 (Figure 3C; ‘MIR-dependent circRNAs’), but had no effect on the production of circTMEM109, circZYMND8 and circSPGAP2C (‘MIR-independent circRNAs’).

**Figure 3.**
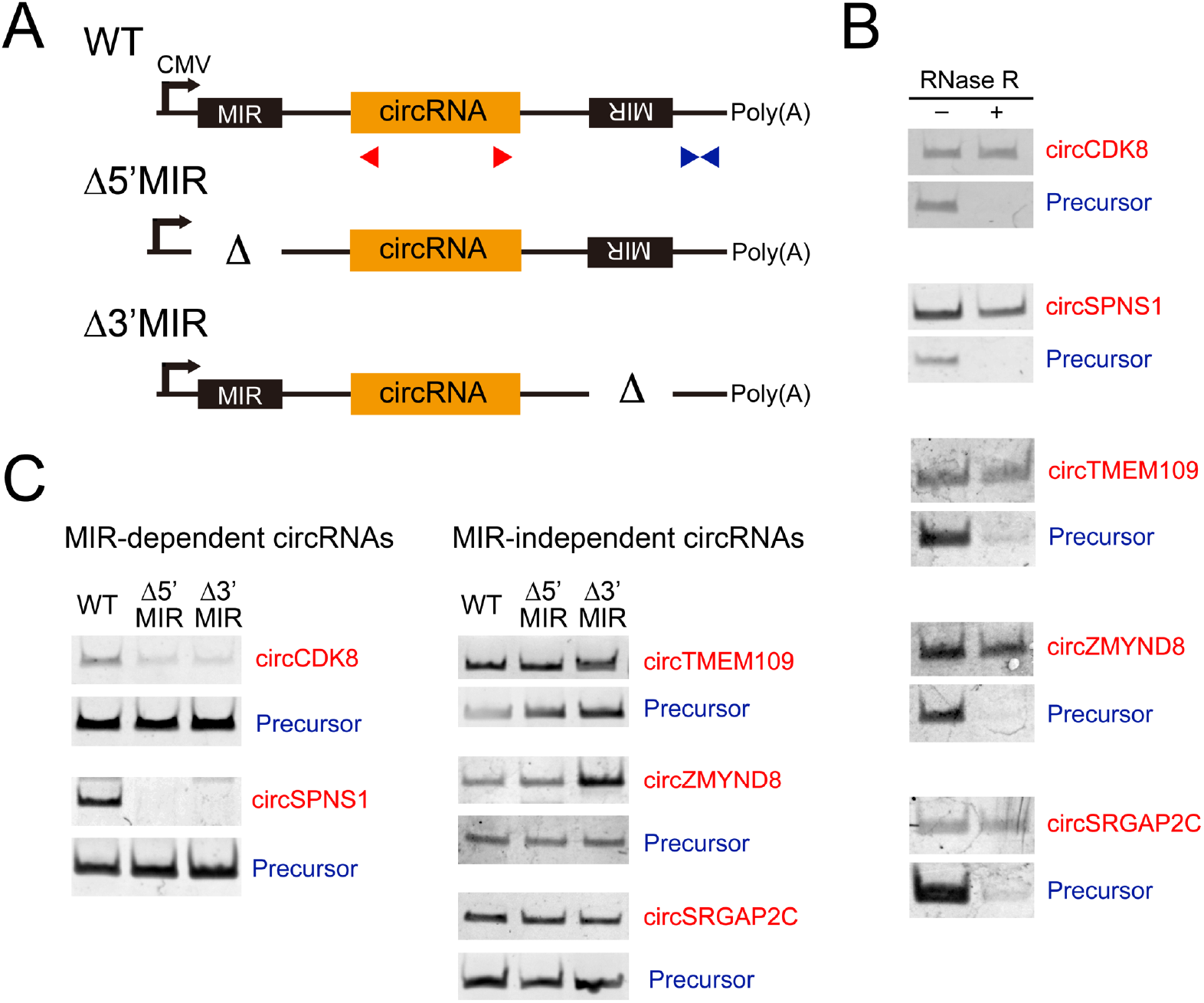
Several Other Human MIR-Dependent circRNAs Were Identified. **(A)** Generalized schematic structures of three kinds of reporters, expressing wild-type (WT) or MIR-deletions (Δ5’MIR, Δ3’MIR) plasmids, which were constructed for five circRNAs. Red and blue triangles indicate the positions of PCR primers to detect these circRNA and the precursors, respectively. **(B)** RT-PCR detections of indicated five circRNAs that were expressed in mouse N2A cells transfected with the plasmids depicted in panel A. Total RNA from the cells was treated with (+) or without (–) RNase R to discriminate the circularized structure from the linear structure. **(C)** Identification of MIR-dependent and MIR-independent circRNAs. RT–PCR analysis of circRNAs and their precursors, which were expressed from plasmids depicted in panel A.

To confirm this in an endogenous setting, CRISPR/Cas9-mediated editing was applied to delete the MIR elements flanking the circCDK8 (Figure S6A). The effective deletion was confirmed by PCR (Figure S6B). Using RT–qPCR, we observed marked inhibition of circularization when either the upstream or downstream MIR sequences were deleted (Figure S6C).

### MIR-MIR Base-Pairing Stability Is Critical for circRNA Generation

MIRs are an older SINE family than *Alus* (~130 vs. ~60 million years ago), and MIR elements have often become truncated during this time (Jurka et al., 1995). We assumed that such truncations would cause instability of the MIR-MIR base-paring required for efficient circRNA production.

To test this hypothesis, we evaluated the integrity of MIRs located around MIR-dependent and MIR-independent circRNAs by comparing them to the MIR consensus sequence using a Smith-Waterman (SW) alignment score (Smith and Waterman, 1981). As predicted, MIRs flanking MIR-dependent circRNAs had markedly higher alignment scores than those of MIR-independent circRNAs (Figure 4A; Figure S5 for the individual score of all MIRs). We also observed that the MIRs of MIR-dependent circRNAs are significantly longer than those of MIR-independent circRNAs (Figure 4B).

**Figure 4.**
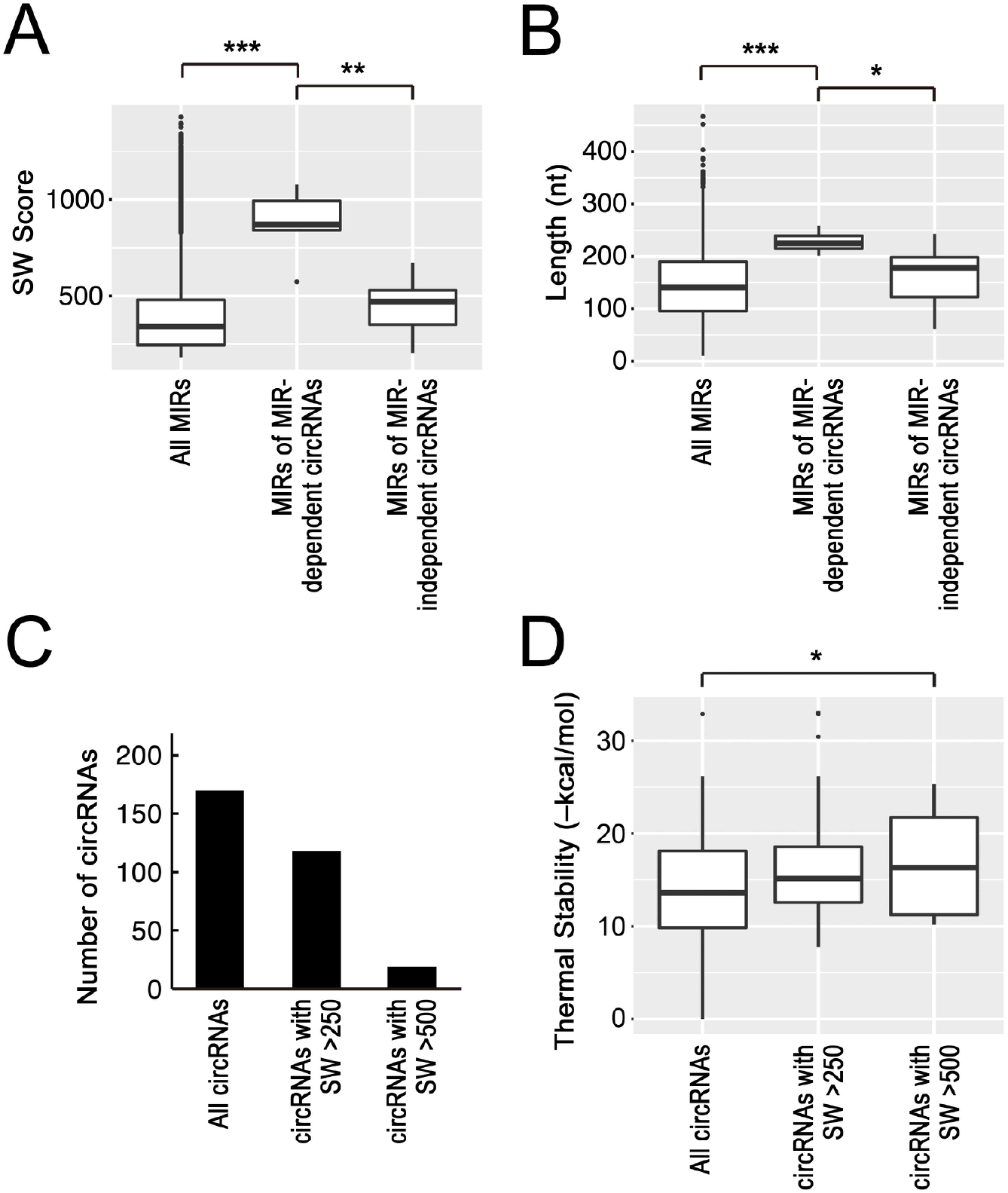
MIR-Dependent circRNAs Likely Depend on Stable Paring between Highly Conserved MIRs. **(A)** The identified MIR-dependent circRNAs have MIRs with near-consensus features. Box plot of SW alignment scores of all reported MIRs, three MIR-dependent circRNAs, and three MIR-independent circRNAs (***P < 0.001; **P < 0.01). See Figure S5 for each SW score of these MIRs. **(B)** The identified MIR-dependent circRNAs have longer MIRs than those of MIR-independent circRNAs. Box plot of the MIR lengths of all reported MIRs, three MIR-dependent circRNAs, and three MIR-independent circRNAs (***P < 0.001; *P < 0.05). **(C)** We identified 170 circRNAs (left bar) that have inverted MIRs within 1000 nt of the circRNA coding exon. Among these circRNAs, 118 circRNAs had MIRs with SW score >250 (middle bar) and 19 circRNAs had MIRs with SW score >500 (right bar). **(D)** The inverted MIRs with higher SW scores have potential to form more stable MIR-MIR base-pairs. Box plot of the corrected thermal stabilities (–kcal/mol) of inverted MIRs from groups of the circRNA shown in panel C, which are calculated by the RNAup program (*P < 0.05).

Using these criteria of the SW alignment score, we estimated the number of MIR-dependent circRNAs from our identified 170 circRNAs with flanking inverted MIRs. We found 118 and 19 circRNAs to have higher SW scores than 250 and 500, respectively (Figure 4C). The high MIR-SW scores (573–1077) seen for the experimentally confirmed MIR-dependent circRNAs (Figure 3C, Figure S5A and the legend) suggest that these 19 circRNAs (>500 MIR-SW scores) are all likely to show MIR-dependent biosynthesis. We calculated the thermal stability of MIR-MIR pairing of these groups of circRNAs, showing that the inverted MIRs with higher SW alignment scores can form more stable MIR-MIR paring (Figure 4D).

Finally, we tried to predict the complementarity of inverted MIRs of all six selected circular RNAs together with three Alu-dependent circRNAs. The MIR-MIR base-parings of MIR-dependent circRNAs are apparently more stable than those of MIR-independent circRNAs, though not so much as those of Alu-dependent circRNAs (Figure 5). Together, we conclude that subset of widespread mammalian circRNAs can be circularized *via* stable RNA pairing of MIRs close to the consensus sequence.

**Figure 5.**
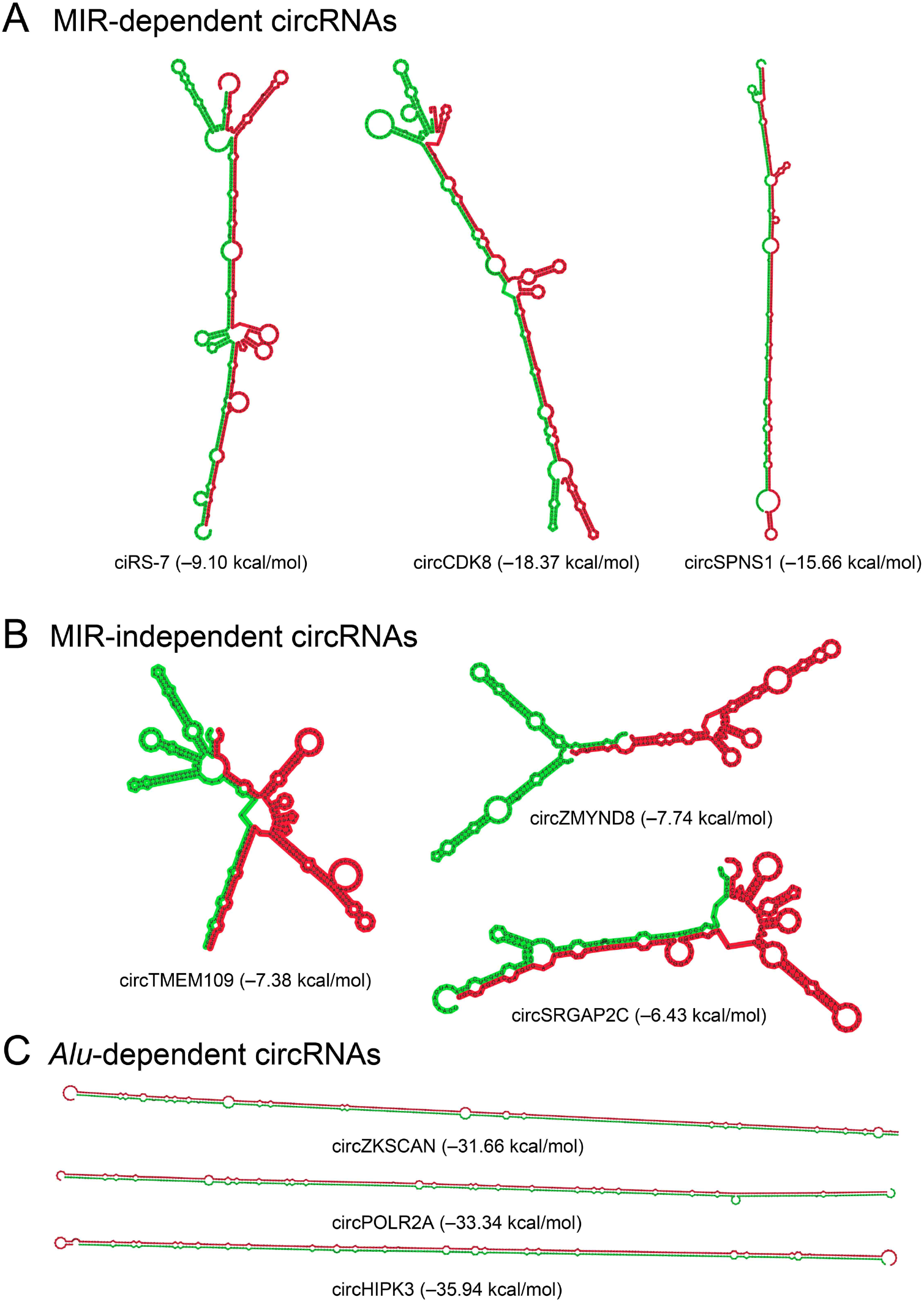
Inverted Sequences Are Less Complementary in MIRs than in *Alus*. RNA secondary structures were predicted by RNAcofold of the Vienna RNA package. The cofold structure of MIR-dependent circRNAs **(A)**, MIR-independent circRNAs **(B)**, and Alu-dependent circRNAs **(C)** are shown. The corrected thermal stability (–kcal/mol) of each MIR was calculated by the RNAup program.

## DISCUSSION

We and others first characterized the circular RNA ciRS-7 as a regulator of a specific miRNA, miR-7 (Hansen et al., 2013; Memczak et al., 2013). Here, we describe the biosynthesis pathway for this important circRNA. Circular ciRS-7 is formed *via* a back-splicing process that is facilitated by flanking inverted sequences derived from MIR, a conserved mammalian SINE. Furthermore, we identified MIR-derived elements that are associated with a number of circRNAs and promote back-splicing and circularization, illustrating a distinct subset of mammalian circRNAs whose formation are driven by MIRs but not by *Alus.*

### Previous Results Are Consistent with the MIR-Dependent Back-Splicing of ciRS-7

Previous ciRS-7 back-splicing assays used reporter plasmids that include the endogenous flanking genomic sequence, 1 kb upstream and 0.2 kb downstream from the ciRS-7 exon (Hansen et al., 2013). This reporter could not produce ciRS-7, consistent with the fact that the inverted MIRs required for back-splicing are located ~3.0 kb and ~0.6 kb away from the 3’ and 5’ splice sites of ciRS-7 exon, respectively.

Recently, the genomic structure around the ciRS-7 precursor has been independently reported (Barrett et al., 2017). In agreement with our analysis, this study showed that the promoter region of ciRS-7 overlaps an upstream long non-coding transcript (LINC00632 locus), suggesting that the ciRS-7 precursor is a long non-coding RNA (Barrett et al., 2017). The BAC clones obtained (185 and 200 kb) cover from the promoter of this non-coding transcript (LINC00632) to far downstream of the ciRS-7 exon and they were found to produce mature ciRS-7. The rationale is that these two BAC clones includes upstream and downstream MIRs that are essential for ciRS-7 generation as we showed here.

### MIR-Dependent Back-Splicing Is Widely Utilized for Mammalian circRNA Biosynthesis

The direct back-splicing pathway is commonly used to generate circRNA in metazoans (Ivanov et al., 2015), however only flanking inverted *Alu* sequences have been identified as back-splicing promoting elements. Indeed *Alus* are major players in the biosynthesis of human circRNAs (Jeck et al., 2013; Zhang et al., 2014) because of their abundance (~10% in human genome) (Price et al., 2004). However, the Alu-SINEs are relatively young elements (~60 million years) and exist only in primates. MIR-SINEs are much older (~130 million years) and globally functional in mammals (Krull et al., 2007), making it likely that MİRs are commonly used in the biosynthesis of mammalian circRNAs.

Due to their age, many MIR sequences have lost their 5’ and 3’ regions, reducing their potential to form complementary duplexes, whereas the more recently emerged *Alu* sequences have remained more complementary to one another. An earlier study found that short complementary sequences (30–40 nt) in the inverted *Alu* elements are required to support efficient back-splicing (Liang and Wilusz, 2014). According to this criterion, the MIR-independent circRNAs may not have to possess complementary sequences longer than 30 nt. Notably, the difference between MIR-dependent and MIR-independent circRNAs was reflected in the predicted secondary structures of their MIR sequences.

### MIR-Deletion Could Be Effective Way To Investigate Function of Targeted circRNA

Our bioinformatics analysis identified 19 circRNAs with conserved flanking MIRs with high complementarity (SW score >500), and these are thus expected to be formed *via* a MIR-dependent pathway. As the representative of these, we verified that circCDK8 and circSPNS1, as well as ciRS-7, were indeed generated through flanking inverted MIR elements.

The host genes of these MIR-dependent circRNAs, circCDK8 and circSPNS1, have important biological roles. CDK8 kinase is a component of the Mediator kinase complex that regulates transcription by RNA Polymerase II (Reviewed in Dannappel et al., 2019). The SPNS1 protein is a putative lysosomal H+-carbohydrate transporter that is required for autophagy (Sasaki et al., 2014; Sasaki et al., 2017; and references therein).

To examine the unknown functions of these MIR-derived circRNAs, it is essential to knock-out the target circRNA while retaining the host gene expression, or the normal spliced mRNA. Now, we can take advantage of the detected MIRs to specifically attenuate circRNA expression using CRISPR/Cas9 editing. Our investigation of the phenotype in the MIR-targeted knockdown cells will shed light on the biological and physiological functions of circCDK8 and circSPNS1.

## EXPERIMENTAL PROCEDURES

### Analysis of RNA-Seq Datasets

We used RNA-Seq datasets from non-neoplastic brain tissue (GSE59612; Gill et al., 2014). Obtained data were aligned to reference human genome hg19 using HISAT2 (Kim et al., 2015) and the aligned sequence reads were assembled by StringTie (Pertea et al., 2015). Repeated sequences were downloaded from the RepeatMasker track in the UCSC table browser (https://genome.ucsc.edu/cgi-bin/hgTables) and analyzed with BEDtools (http://bedtools.readthedocs.io).

The box plots were constructed using the R/Bioconductor package (http://www.bioconductor.org). For statistical comparisons of groups, Wilcoxon rank-sum tests were used to calculate P-values.

Possible RNA pairings between inverted MIRs were predicted using the RNAcofold program from the Vienna RNA package (Lorenz et al., 2011). Thermodynamic stability of the base-paired MIRs was calculated with the RNAup program in the Vienna RNA package using ‘–b’ option to include the probability of unpaired regions (Lorenz et al., 2011). Exceptionally weak MIR-MIR base-pairings that the RNAup program could not calculate their thermodynamic stabilities were set to zero values (‘All circRNAs’ in Fig. 4D).

### RT–PCR Assays

RT–PCR analysis was performed essentially as previously described (Yoshimoto et al., 2017). Human cerebral cortex total RNA was purchased from Clontech (CLN 636561). From culture cells, total RNA was isolated with a Nucleospin RNA kit (Machery Nagel) according to the manufacturer’s instructions. ‘On-column’ DNase I digestion was performed to remove contaminated DNA. Purified RNA was reverse-transcribed using PrimeScript II (Takara Bio) with oligo-dT and random primers, and cDNA was amplified by PCR (20–30 cycles) with Ex-Taq (Takara Bio) and specific primers (Table S1). PCR-amplified products were analyzed by 8% polyacrylamide gel electrophoresis.

For nested PCR, the first PCR reaction mixture (30 cycles) was purified with a PCR cleanup column (Takara Bio) and the eluate was used for the second PCR reaction (30 cycles). The purified RNase R (1 μg) was added to digest 1 μg of total RNA to remove linear RNA in a 20 μL reaction mixture at 30°C for 30 min as described previously (Suzuki et al., 2006).

To perform quantitative PCR (qPCR), a real-time PCR instrument (Eco Real-Time PCR System, Illumina) was used with the same primers (Table S1) that were used in the regular PCR. For the qPCR reactions, KAPA Taq PCR kit was used according to the manufacturer’s protocol (Kapa Biosystems).

For the RT-qPCR analysis in Figure 2E, DNase-treated RNA (1 μg) was reverse-transcribed using a Superscript VILO cDNA Synthesis Kit (Thermo Fisher Scientific) according to the manufacturer’s instructions. Obtained cDNA was mixed with SYBR Green I Master (Roche Molecular Systems) and analyzed on a real-time PCR instrument (LightCycler 480, Roche Molecular Systems) according to the manufacturer’s protocol.

### Northern Blot Analyses

Total RNAs (5 μg each) were separated by electrophoresis on 1% agarose containing 3% formaldehyde, rinsed twice in distilled water for 10 min, denatured in 7.5 mM NaOH for 20 min, neutralized in 20× SSC containing 3 M NaCl and 0.3 M sodium citrate (pH 7.0) for 20 min, and blotted overnight on a nylon membrane (RPN82B, GE Healthcare Life Sciences) followed by UV irradiation at 254 nm with 120 mJ/cm^2^ (CL-1000, Funakoshi). DIG-labeled probes were hybridized in PerfectHyb solution (HYB-101, Toyobo) overnight at 68°C, and hybridized probes were detected by alkaline phosphatase-conjugated anti-DIG antibodies (#11093274910, Roche) and CDP-star (#CDP-RO, Roche). The chemiluminescence signals were observed by imaging analyzer (ImageQuant LAS 500, GE Healthcare Life Sciences).

### Antisense Oligoribonucleotide-Mediated Splicing Repression

Antisense 2’-O-methyl-modified phosphorothioate oligoribonucleotides (ASOs) were purchased from Integrated DNA Technologies (Table S1). These chemically modified ASOs were electroporated into SH-SY5Y cells (Gene Pulser Xcell, Bio-Rad). Fully confluent SH-SY5Y cells grown on 10 cm plate in D-MEM/Ham’s-F12 medium (Fujifilm Wako) were trypsinized and the washed cell pellets were suspended in 1 mL OPTI-MEM medium (Thermo Fisher Scientific). This cell suspension (0.2 mL) plus ASO (2.5–10 μM final concentration) were transferred into 0.4 cm cuvette (BEX) and electroporated at 200 V for 20 ms square-wave pulses. After electroporation, cell suspensions were transferred into 6-well plates supplemented with 1.8 mL D-MEM/Ham’s-F12 that was cultured for 24 h before RT–PCR analysis.

### Construction and Expression of circRNA-Reporter Plasmids

The expression plasmids for ciRS-7, circCDK8, circTMEM109, circZMYNDR8, and circSRGAP2C were constructed by subcloning the PCR-amplified fragments into FLAG-pcDNA3 vector using BamHI and XhoI sites (Yoshimoto et al., 2009). The expression plasmids for circSPNS1 was constructed by subcloning the PCR-amplified fragments into FLAG-pcDNA3 plasmid using an In-Fusion HD cloning kit (Takara Bio) according to the manufacturer’s protocol. The deletion of MIR element of these expression plasmids were made by conventional extension PCR method.

To establish HEK293T cells stably expressing ciRS-7, PCR-amplified a ciRS-7 fragment was subcloned into pcDNA5 FRT-TO vector (Thermo Fisher Scientific) using BamHI and XhoI sites, and transfected into Flp-In T-REx 293 cells (Thermo Fisher Scientific) along with Flp recombinase vector pOG44. The transfected cells were treated with 1 μg/mL DOX for 24 h and total RNA was prepared for the RT–PCR analysis. To inhibit the splicing reaction, 100 ng/mL SSA was added 18 h after DOX treatment.

Mouse N2A cells were transiently transfected with these expression plasmids using Lipofectamine 2000 (Thermo Fischer Scientific) according to the manufacturer’s instructions. The transfected cells were incubated for 24 h and total RNA was prepared for RT–PCR analysis.

### Targeted Genomic Deletion of the Splice Sites in the ciRS-7 Locus and of MIRs in the circCDK8 Locus

Cas9 RNP was introduced with the Alt-R CRISPR-Cas9 System (Integrated DNA Technologies) according to the manufacturer’s instructions. The gRNAs in Figures S4A, S6A and primer sequences are listed in Table S1.

HEK293 cells were seeded into a 96-well plate at a density of 4×10^4^ cells/well with DMEM/F-12 medium (Fujifilm Wako Pure Chemical) containing 10% fetal bovine serum (FBS, Sigma-Aldrich) and 0.5% Gibco penicillin-streptomycin mixture (Thermo Fisher Scientific). The cells were transiently transfected with 0.75 pmol of Cas9 RNP using 1.2 μL of Lipofectamine RNAiMAX (Life Technologies) diluted up to 50 μL of OPTI-MEM (Thermo Fisher Scientific). After a 20 min incubation at room temperature, the transfection solution was added dropwise to the cells. At 48 h post-transfection, the cells were trypsinized and filtered through a 40 μm Corning cell strainer (Thermo Fisher Scientific), seeded onto a 10-cm culture plate, and grown for 2 weeks. Each colony (~96 clones per construct) were picked, trypsinized, and seeded onto 96-well plate with two replicates. Genomic DNA was extracted and PCR was performed to verify the deleted region.

### Targeted Genomic Deletion of MIRs in ciRS-7 Locus

The gRNAs (Figure 2C) are listed in Table S1. The annealed gRNAs were subcloned into the BpiI-cleaved (Thermo Fisher Scientific) site of a modified version of CRSPR vector px458 (#43138, Addgene) without the ITR element that expresses Cas9/EGFP (Højland Knudsen et al., 2018). To increase the efficiency and accuracy of the CRISPR targeting site (MIR elements), single-stranded donor oligodeoxyribonucleotides (ssODN) acting as repair templates were used (Table S1).

SH-SY5Y cells were seeded in six-well plates and grown to 70% confluency (0.8×10^6^ cells/well) in DMEM/F-12 medium containing GlutaMax, 10% FBS, and 0.5% Gibco penicillin-streptomycin mixture (all reagents from Thermo Fisher Scientific). The cells were transiently transfected with 2.5 μg DNA (1 μg each of two gRNA vectors and 0.5 μg ssODN) using Lipofectamine 3000 (Thermo Fisher Scientific) according to the manufacturer’s instructions. After 15 min incubation at room temperature, the transfection solution was added dropwise to the cells. The medium was replaced with fresh growth medium at 12 h post-transfection.

At 48 h post-transfection, the cells were prepared for fluorescent-activated cell sorting (FACS) by trypsinization, washing, and re-suspension in PBS with 2% FBS. The cells were FACS sorted in a BD FACSAria III (BD Biosciences) based on their viability and GFP signal quality into 96-well plates (single cell per well) containing 150 μl of conditioned growth medium (30% used and 70% fresh media). Three weeks after the FACS sorting, the grown colonies were passaged into duplicate 96-well plates and one set was used for DNA preparation and genomic PCR with flanking primers to detect the deleted homozygote clones. The verified deleted genomic clones and the appropriate controls were expanded and total RNA was prepared for RT-qPCR analysis (see above).

## ACKNOWLEDGEMENT

We are grateful to Drs. Kinji Ohno and Hitomi Tsuiji for SH-SY5Y cells and N2A cells, respectively. Spliceostatin A (SSA) was a generous gift from Dr. Minoru Yoshida. We thank Drs. Anne Nielsen and Shinichi Nakagawa for critical reading of the manuscript, Dr. Makoto K. Shimada for helpful suggestions for bioinformatics analysis, Taiwa Komatsu and Mayuko Tanahashi for providing instruction of the gene editing techniques, and members of our laboratory for constructive discussion.

R.Y. was supported by a Research Grant from the Hori Sciences and Arts Foundation, a Research Grant from the Nitto Foundation, a Science Research Promotion Fund from the Promotion and Mutual Aid Corporation for Private Schools of Japan (PMAC), and Grants-in-Aid for Scientific Research (C) (JP18K05563). A.M. was partly supported by Grants-in-Aid for Scientific Research (B) (JP16H04705) and Grants-in-Aid for Challenging Exploratory Research (JP16K14659) from the Japan Society for the Promotion of Science (JSPS).

## AUTHOR CONTRIBUTIONS

R.Y. and A.M. conceived and designed the experiments; R.Y. performed most of the experiments and analyses, organized the data and drafted the manuscript; K.R. performed CRISPR/Cas9-mediated genomic deletion of the MIR elements; T.H. and J.K. provided useful information and revised the manuscript; R.Y. and A.M. analyzed the data and edited the manuscript. A.M. coordinated and supervised the whole project. All authors read, corrected and approved the final manuscript.

## Notes

#### Summary of Updates

Text was revised and reorganized; Figures were replaced and rearranged; Supplemental files (including Table S1) were updated.

