## Supplemental Figures S1-S6 for "Biosynthesis of Circular RNA ciRS-7/CDR1as Is Mediated by Mammalian-Wide Interspersed Repeats (MIRs)"

### Contents

**Figure S1**

**Figure S2**

**Figure S3**

**Figure S4**

**Figure S5**

**Figure S6**

**Table S1**
(Uploaded in a separate Excel file)

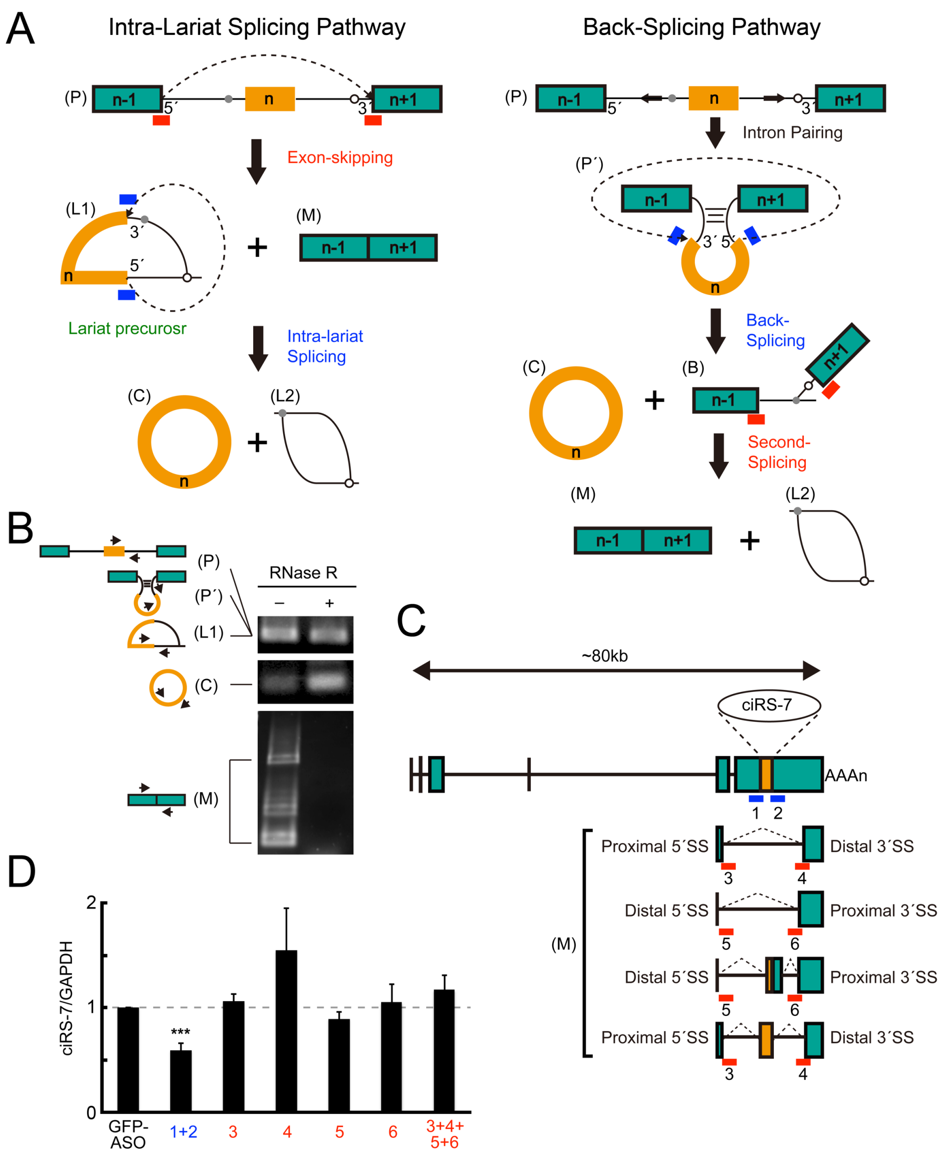


**Figure S1. Evidence for 'Back-splicing' rather than 'Intra-lariat splicing' to produce circular ciRS-7**
**(A)** Two proposed pathways for circRNA biosynthesis, which can be distinguished by blocking of specific splicing events using antisense oligoribonucleotides (ASOs). The ASOs 1+2 (blue bars) targeting the flanking 5′ and 3′ splice sites of ciRS-7 exon can prevent circular ciRS-7 generation *via* both pathways (used as the positive control). The ASOs 3–6 (red bars) targeting alternative 5′ and 3′ splice sites outside of ciRS-7 exon can prevent the exon-skipping and the following circular ciRS-7 generation in the 'Intra-lariat pathway' (left panel). However, these same ASOs 3–6 cannot interfere with the back-splicing event, and thus allow circular ciRS-7 generation in the 'Back-splicing pathway' (right panel).
**(B)** Detection of final closed circular form of ciRS-7 (C) together with linear form of several final spliced products (M) due to multiple alternative 5′ and 3′ splice sites (see panel C). Total RNA from human cerebral cortex was treated with (+) or without (–) RNase R to distinguish opened linear RNA from closed RNA. The purified RNA was analyzed by RT–PCR with indicated primers (arrowheads).
**(C)** Schematic representation of the ciRS-7 precursor and the alternatively spliced isoforms (see panel B and Figure S2). Bars indicate ASOs (1–6) to block specific alternative splicing event.
**(D)** The quantified data of splicing prevention using ASOs (1–6; see panel C). SH-SY5Y cells were electroporated with each ASO; the ASOs 1+2 significantly repressed ciRS-7 production (blue), whereas none of the ASOs 3–6 inhibited ciRS-7 production (red). RT–PCR analysis was performed with ciRS-7 primers and the control GAPDH primers. The detected ciRS-7 expression levels were normalized to the control expression level of GAPDH (ciRS-7/GAPDH) and plotted as ratios to the value of control cells treated with GFP-ASO. Means ± standard deviation (SD) are given for three independent experiments (***$P < 0.001$).

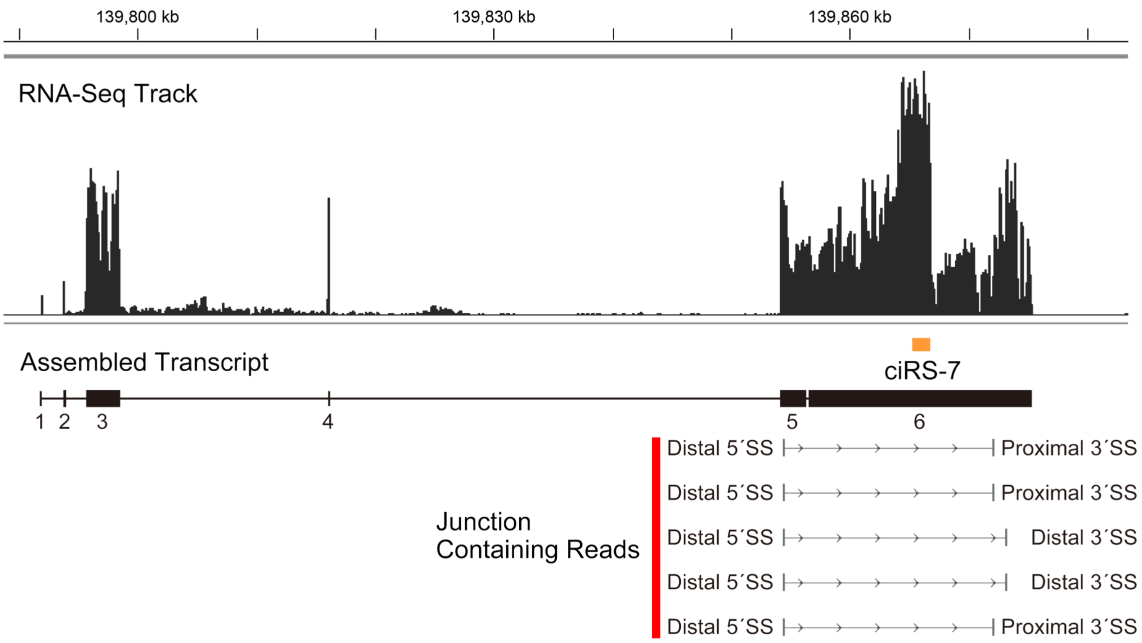


**Figure S2. Alternative splicing events skip the ciRS-7 exon.**
Modified screeenshot of human brain RNA-Seq data (GSE59612) mapped on the the human hg19 genome sequence is indicating the assembled ciRS-7 precursor transcript. The junction containing reads revealed the active alternative 5′ and 3′ splice sites within the exons 5 and 6.

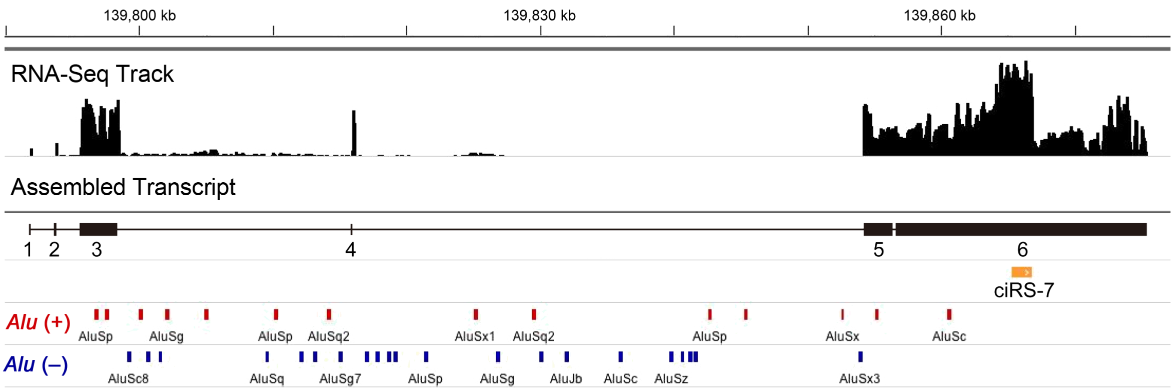


**Figure S3. The ciRS-7 exon does not have flanking inverted *Alu* elements.**
The locations of the human *Alu* elements are shown together with the assembled ciRS-7 precursor tarnscript (see Fig. S2 above). The *Alu* elements (bars) were extracted from UCSC Repeat Masker database (+ and – are the same and opposite strand of the ciRS-7 exon, respectively).

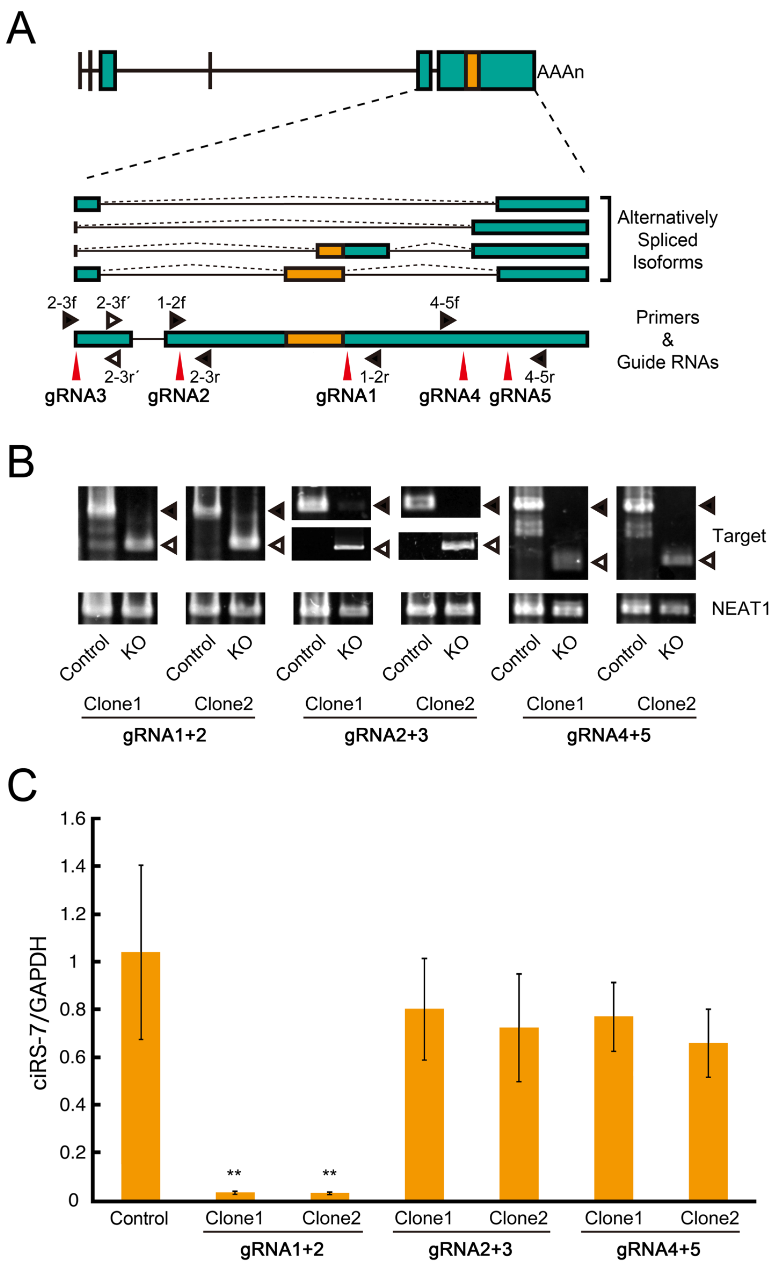


**Figure S4. CRISPR/Cas9-mediated deletion of the flanking alternative splice sites of ciRS-7 supports 'Back-splicing pathway' to generate circular ciRS-7.**

**(A)** Schematic structures of the ciRS-7 precursor and its alternatively spliced isoforms (orange box indicates ciRS-7 exon). The positions of the guide RNAs (gRNA1–gRNA5) targetting the alternative splice sites, PCR primers for detecting deleted sites (filled triangles), and those for detecting un-deleted sites (open triangeles) are also indicated.
**(B)** The targeted ciRS-7 genomic deletions in HEK-293 cells were verified by genomic PCR. The indicated three pairs of gRNAs were used to delete the alternative splice sites. PCR primers indicated in panel A were used for detecting deleted sites (open triangles) and non-deleted sites (filled triangles). PCR primers for the *NEAT1* gene were used as control.
**(C)** The deletion of either the alternative 5′ splice sites (between gRNA2 and gRNA3) or the 3′ splice sites (between gRNA4 and gRNA5) outside of ciRS-7 exon barely prevented the generation of ciRS-7. The ciRS-7 production was analyzed by quantitative RT–PCR with ciRS-7 primers and the control GAPDH primers. The ciRS-7 expression levels were normalized to the control expression level of GAPDH (ciRS-7/GAPDH). The results were plotted as ratios to the value of control wild-type cells. Means ± standard deviation (SD) are given for three independent experiments (**P < 0.01).

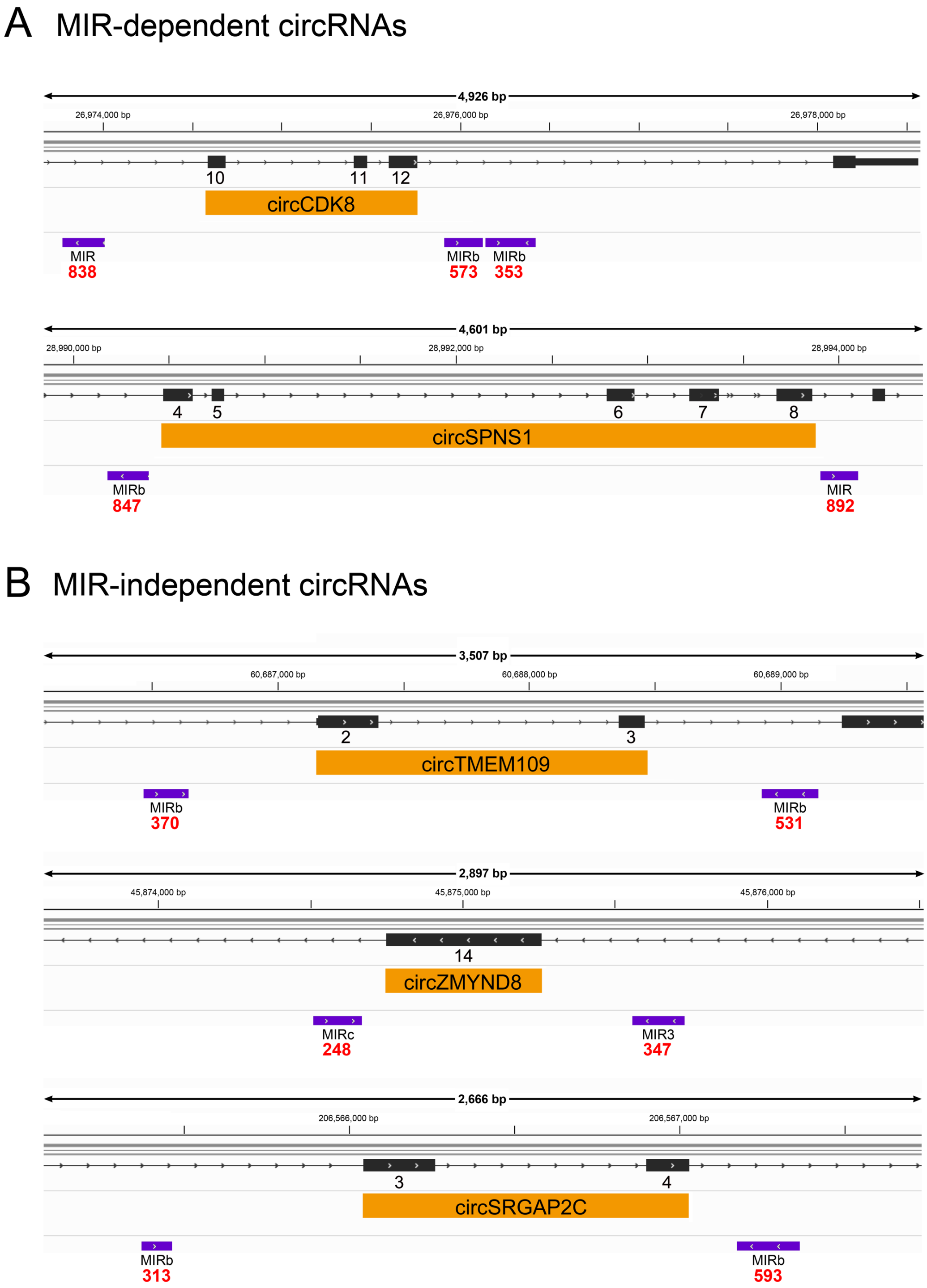


**Figure S5. Identified another human circRNAs revealed different MIR-dependency.**
The loci of two MIR-dependent circRNAs **(A)** and three MIR-independent circRNAs **(B)** are shown. The positions of the host genes (with red numbered exons), circRNA exons (orange boxes), and the identified inverted MIR elements (purple bars) are indicated. Red numbers indicate the Smith-Waterman (SW) alignment scores of these MIR elements. The SW alignment scores of MIRs in MIR-dependent human ciRS-7 (see Fig. 1A) are 945 (upstream) and 1077 (downstream).

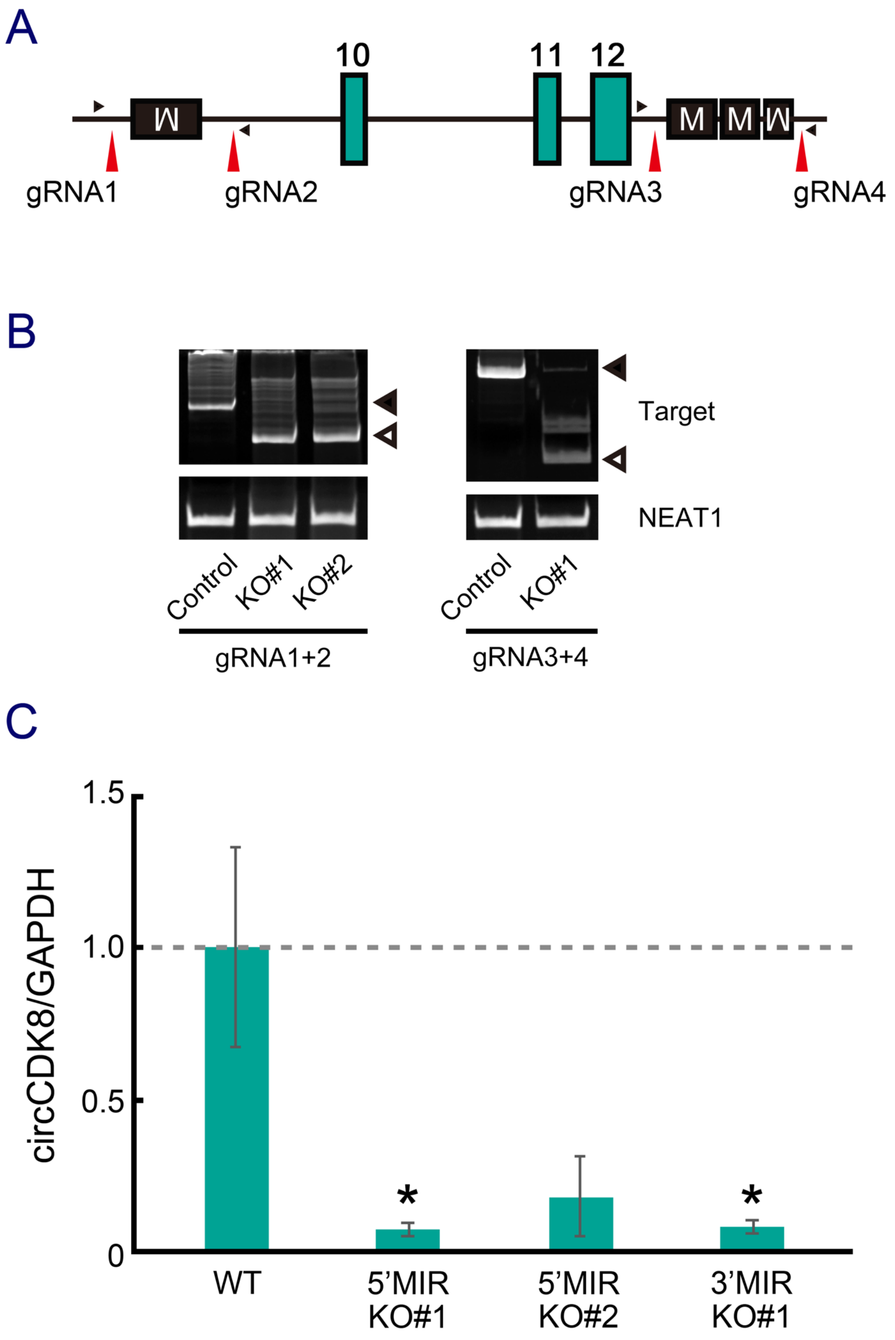


**Figure S6. CRISPR/Cas9-mediated deletion of either upstream or downstream MIRs aborts the production of circCDK8**

**(A)** Schematic genomic structure of circCDK8 locus with the flanking inverted MIR sequences. The positions of guide RNAs (gRNA1–gRNA4) to delete each MIR element are indicated with red vertical arrowheads. PCR primers for detecting deleted sites are indicated with filled triangles.

**(B)** The genomic deletions of the flanking MIR elements in HEK293 cells were verified by genomic PCR. The indicated two pairs of gRNAs were used to delete the 5′ and 3′ MIR elements. PCR primers indicated in panel A were used for detecting the MIR-deleted sites (open triangles in KO#1 and KO#2) and non-deleted sites (filled triangles in Control).

**(C)** The effects of the MIR deletions (5′MIR KO and 3′MIR KO) on the circCDK8 production was quantified by qRT–PCR. The ciRS-7 expression levels were normalized to the control expression level of GAPDH (circCDK8/GAPDH). Values are relative to the value of undeleted parental cells (WT). Means ± standard deviation (SD) are given for three independent experiments (*P < 0.05).
